# Laryngeal and swallow dysregulation following acute cervical spinal cord injury

**DOI:** 10.1101/2021.09.21.457706

**Authors:** Teresa Pitts, Kimberly E. Iceman, Alyssa Huff, M. Nicholas Musselwhite, Michael L. Frazure, Kellyanna C. Young, Clinton L. Greene, Dena R. Howland

## Abstract

Proper function of the larynx is vital to airway protection, including swallow. While the swallow reflex is controlled by the brainstem, patients with cervical spinal cord injuries (cSCI) are likely at increased risk of disordered swallow (dysphagia) and pneumonia, and the underlying mechanisms are unknown. We aimed to determine if acute spinal cord injury would disrupt swallow function in animal models. We hypothesized that 1) loss of descending efferent information to the diaphragm would affect swallow and breathing differently, and that 2) loss of ascending spinal afferent information would alter central swallow regulation to change motor drive to the upper airway. We recorded amplitudes of laryngeal and inspiratory muscle electromyograms (EMGs), submental and pharyngeal muscle EMGs, and cardiorespiratory measures in freely breathing pentobarbital-anesthetized cats and rats. First, we assessed the effect of a lateral hemisection at the second cervical level (C2) in cats during breathing. Posterior cricoarytenoid (laryngeal abductor) EMG activity during inspiration increased nearly two-fold, indicating that inspiratory laryngeal drive increased following cSCI. Ipsilateral to the injury, the crural diaphragm EMG was significantly reduced during breathing (62 ± 25 percent change post-injury), but no animal had a complete termination of all activity; 75% of animals had an increase in contralateral diaphragm recruitment after cSCI, but this did not reach significance. Next, we assessed the effect of C2 lateral hemisection in cats during swallow. The thyroarytenoid (laryngeal adductor) and thyropharyngeus (pharyngeal constrictor) both increased EMG activity during swallow, indicating increased upper airway drive during swallow following cSCI. There was no change in the number of swallows stimulated per trial. We also found that diaphragm activity during swallow (schluckatmung) was bilaterally suppressed after lateral C2 hemisection, which was unexpected because this injury did not suppress contralateral diaphragm activity during breathing. Swallow-breathing coordination was also affected by cSCI, with more post-injury swallows occurring during early expiration. Finally, because we wanted to determine if the chest wall is a major source of feedback for laryngeal regulation, we performed T1 total transections in rats. As in the cat C2 lateral hemisection, a similar increase in inspiratory laryngeal activity (posterior cricoarytenoid) was the first feature noted after rat T1 complete spinal cord transection. In contrast to the cat C2 lateral hemisection, diaphragmatic respiratory drive increased after T1 transection in every rat (215 ± 63 percent change), and this effect was significant. Overall, we found that spinal cord injury alters laryngeal drive during swallow and breathing, and alters swallow-related diaphragm activity. Our results show behavior-specific effects, suggesting that swallow may be more affected than breathing is by cSCI, and emphasizing the need for additional studies on laryngeal function during breathing and swallow after spinal cord injury.

## Introduction

The larynx and the upper esophageal sphincter act in concert to control the flow of air and food into the lungs and esophagus respectively (Pitts, Rose et al. 2013). Laryngeal dysfunction, whether central or peripheral in origin, leads to a wealth of behavioral deficits including dysphonia (disorder of voice) (Barton 1979, Allegretto, Morrison et al. 2003, Cukier-Blaj, Bewley et al. 2008, Neri, Castiello et al. 2011), dystussia (disorder of cough) (Fontana, Pantaleo et al. 1998, Fontana, Pantaleo et al. 1999, Smith Hammond, Goldstein et al. 2001, Smith Hammond, Goldstein et al. 2009), and dysphagia (disorder of swallow) (Ali, Wallace et al. 1996, Rosenbek, Robbins et al. 1996, Bath, Bath et al. 1999, Kendall and Leonard 2001, Daggett, Logemann et al. 2006). While all these symptoms occur after spinal cord injury, dysphagia has received the least attention.

Swallow has 3 phases: oral, pharyngeal, and esophageal (Bosma 1957, Doty 1968, Miller 1982, Bieger 2001). The pharyngeal phase, which is the focus of this manuscript, directs food/liquid past the larynx and through the upper esophageal sphincter, using a highly regulated series of coordinated, bilateral muscle contractions over ∼500 ms (Doty and Bosma 1956, Doty 1968, Thexton, Crompton et al. 2007, German, Crompton et al. 2009). There are two distinct forces on the bolus to propel movement: positive pressure from oropharyngeal contraction and negative pressure from activation of the diaphragm and other inspiratory chest wall muscles (McConnel 1988). Full closure of the laryngeal opening is required to sustain the pressure necessary for effective movement of the bolus into the esophagus, and a leak in this process significantly increases aspiration risk (Ding, Fung et al. 2015).

While dysphagia can have a myriad of origins (Gordon, Hewer et al. 1987, Buchholz 1994, McConnel and O’Connor 1994, Coates and Bakheit 1997, Miller 2008, Sue Eisenstadt 2010), the outcome is often the same: penetration (food/liquid entering the larynx) or aspiration (food/liquid below the larynx in the trachea/lungs) (Rosenbek, Robbins et al. 1996, Robbins, Coyle et al. 1999, Aviv, Spitzer et al. 2002, Daggett, Logemann et al. 2006, Cvejic, Harding et al. 2011), which significantly increases the risk of pneumonia (Mandell and Niederman 2019). While the swallow reflex is brainstem-mediated and the upper airway muscles are controlled by cranial nerves, some studies suggest that spinal cord injury may cause dysphagia (Chaw, Shem et al. 2012, McRae 2015, McRae, Smith et al. 2019). Clinical work in patients with cervical spinal cord injuries (cSCI) has shown mixed results regarding aspiration risk. Reported incidences vary considerably (16-75% prevalence) and until recently were only evaluated retrospectively (Kirshblum, Johnston et al., Wolf and Meiners, Abel, Ruf et al., Brady, Miserendino et al., Shem, Castillo et al., Seidl, Nusser-Müller-Busch et al., Shem, Castillo et al., Shin, Yoo et al., Chaw, Shem et al., Shem, Castillo et al., Shem, Castillo et al., Lee, Gross et al., Hayashi, Fujiwara et al., Ihalainen, Rinta-Kiikka et al., Ihalainen, Rinta-Kiikka et al.). In a 2017 study (Ihalainen, Rinta-Kiikka et al. 2017) in which 37 cSCI patients were prospectively evaluated, 74% penetrated or aspirated when evaluated with videofluoroscopy (an x-ray of swallow in which the bolus is laced with barium to observe function). Of the patients with severe dysphagia, 73% had silent aspiration, which means there were no outward signs (cough or throat clear) and suggests the likelihood of significant risk of underdiagnosis. When dysphagia is diagnosed, treatment options are extremely limited. Feeding tube placement is most common. Although this effectively bypasses swallow, decreasing pneumonia risk by ∼40% (Ramczykowski, Gruning et al. 2012), it negatively affects quality of life and does not promote recovery. Despite recent studies that do report dysphagia in cSCI patients (Frankel, Coll et al. 1998, Hayashi, Fujiwara et al. 2017, Ihalainen, Rinta-Kiikka et al. 2017, Richard-Denis, Erhmann Feldman et al. 2017), none identify a mechanism for dysfunction.

For these reasons, we aimed to determine if an acute lateral hemisection at the second cervical level (C2) in cats would disrupt swallow function due to loss of descending efferent connections to the diaphragm and/or loss of the ascending afferent information necessary for laryngeal regulation and appropriate swallow pattern generation. Our first hypothesis was that loss of descending efferent information to the diaphragm would affect swallow and breathing differently. We based this partly on clinical reports that patients with a loss of negative intra-thoracic pressure had dysphagia, yet showed no complimentary disorders of breathing (McConnel and O’Connor 1994, Bassotti, Germani et al. 1998, Bulow, Olsson et al. 2001, Hoffman, Mielens et al. 2012). Our second hypothesis was that loss of ascending spinal afferent information would alter the excitability of the swallow pattern generator, and thus result in changes in motor drive to the upper airway. While the cat model offers a robust and nuanced study of swallow, the majority of work in the field is completed in the rat. We also performed rat T1 transection experiments to complement our hypothesis testing and to offer a cross-species verification of the effects of spinal cord injury on laryngeal drive.

## Methods

### Study Design

All experiments were approved by the Institutional Animal Care and Use committee of the University of Louisville and conducted in accordance with the American Physiological Society’s Animal Care Guidelines (Drummond 2009). The cat experiments were performed using 4 purpose-bred sexually-intact adults (1-2 years of age) — 2 females (3.3 & 4.0kg) and 2 males (4.4 & 4.8kg). The rat experiments used 5 female Sprague Dawley ex-breeders with an average weight of 0.46 ± 0.05 kg. The number of animals was predetermined before initiation of the experiments and was based on pilot studies not included in this manuscript. Those preliminary studies indicated a robust increase in laryngeal activity after spinal cord injury. An *a priori* power analysis was conducted to test the difference between two independent group means using a two-tailed test, an effect size (Cohen’s *d*= .80), and an alpha of 0.05. For the cat experiments, the expected mean difference was 75 with a SD of 30, and the result showed a total sample of 4 with an actual power of 0.92. For the rat experiments, the expected mean difference was 70 with a population SD of 34, and the result showed a total sample of 5 with an actual power of 0.93.

Our objectives were to determine any changes in 1) motor drive during swallow and breathing following C2 lateral hemisection in the cat, and 2) laryngeal drive during breathing following T1 complete transection in the rat. The primary outcome measures are amplitudes of laryngeal and inspiratory muscle electromyograms (EMGs). We also collected the following additional measures: EMGs of both submental and pharyngeal muscles, arterial blood pressure, arterial blood gas pressures, end-tidal CO_2_, body temperature, and heart and respiratory rates. Paired Student t-tests or ANOVA were performed when appropriate.

The breathing phases were defined as follows: a) inspiration was defined as the onset to the peak of diaphragm activity; b) E1 (i.e. post-I or yield) was defined as the peak of diaphragm activity to the end of the thyroarytenoid burst; c) E2 was defined as the end of E1 to the beginning of inspiration. Swallow-breathing coordination was assessed by noting the phase of breathing in which swallow was initiated. To quantify the phases in which swallow occurred, we assigned swallows during inspiration, E1, and E2 phases to the numbers 1, 2, and 3 respectively, as we have previously published (Pitts, Rose et al. 2015, Reed, English et al. 2019). To assess swallow-breathing coordination, a Wilcoxon Signed Rank Test was used. An assigned coding system was used for the breathing phase in which the swallow occurred: inspiration as “1”; early expiration (i.e. yield or E1) (Huff, Reed et al. 2020) as “2”; and mid/late-expiration as “3”. For all tests a difference was considered significant if the *p*-value was less than or equal to 0.05.

### Electrophysiology recording

All muscle activity was recorded via electromyography (EMG) using bipolar insulated fine wire electrodes (A-M Systems stainless steel #791050) according to the technique of Basmajian and Stecko (1962). The signals were amplified (Grass P511 AC Amplifiers, Natus Neurology), recorded at a 10K sampling rate (1401 Power3 + ADC16 Expansion, Cambridge Electronic Design) and analyzed using Spike 2 (v7, Cambridge Electronic Design). Moving averages of EMGs were integrated with a 20 ms time constant, and exported to CorelDRAW 2020 (v22.1.1.523) for figure creation.

### Cat Model

Animals were initially anesthetized with sevoflurane (3-5%) and then transitioned to sodium pentobarbital (35-40 mg/kg i.v.); supplementary sodium pentobarbital doses were administered as needed (1-3 mg/kg i.v.). A dose of atropine sulfate (0.1-0.2 mg/kg, i.v.) was given at the beginning of each experiment to reduce airway secretions. Cannulas were placed in the femoral artery, femoral vein, and trachea. Arterial blood pressure (BPM-832 Pressure Monitor, Charles Ward Electronics) and end-tidal CO_2_ (Capnomac Ultima, Datex Engstrom) were continuously monitored. Body temperature was monitored and maintained at 37.5 ± 0.5 °C (Homeothermic Monitor, Harvard Apparatus). Arterial blood samples were periodically removed for blood gas analysis (VetScan I-STAT & CG4+ cartridges, Abaxis). PO_2_ was maintained using air mixtures with enriched oxygen (25-60%) to achieve values above 100 mm Hg if needed (RMA-14-SSV, Dwyer Instruments, Inc & GSM-3 Gas Mixer, Charles Ward Electronics). Anesthetic level was evaluated using end-tidal and pCO_2_, as well as reflex activity (jaw tone, pull back, and ocular reflexes).

Swallow was stimulated by infusion of 3cc of water into the oropharynx and evaluated using the following muscles: mylohyoid, geniohyoid, thyrohyoid, thyropharyngeus, thyroarytenoid, cricopharyngeus, parasternal, and crural diaphragms. These muscles span the actions during the pharyngeal phase of swallow: a) mylohyoid, geniohyoid and thyrohyoid for hyolaryngeal elevation; b) thyropharyngeus for inferior pharyngeal constriction; c) cricopharyngeus for upper esophageal sphincter regulation; d) thyroarytenoid for laryngeal adduction; and e) parasternal, costal diaphragm, and bilateral crural diaphragm schluckatmung activity (i.e. inspiratory activity during swallow which creates a negative intra-thoracic pressure) (Pitts, Rose et al. 2013, Spearman, Poliacek et al. 2014, Pitts, Gayagoy et al. 2015, Pitts, Rose et al. 2015). Breathing motor patterns were evaluated using the posterior cricoarytenoid, thyroarytenoid, and bilateral crural diaphragm. Respiratory phase duration measured with activity from inspiratory and expiratory laryngeal muscles is described in Figure 1.

**Figure 1.**
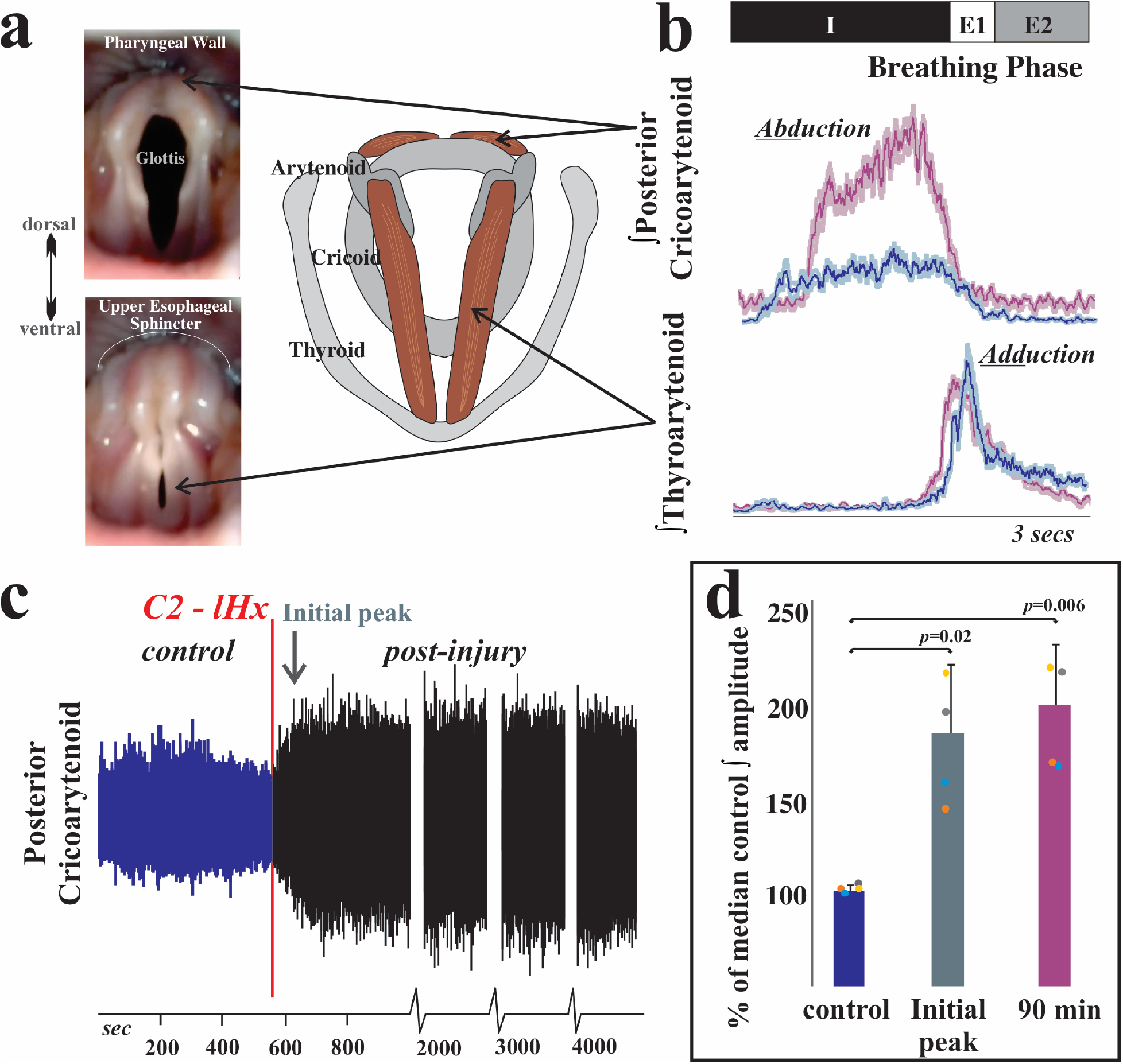
Inspiratory laryngeal drive increases following cervical spinal cord injury. a) Endoscopic laryngeal images of a spontaneously breathing, spinally intact cat (isoflurane anesthesia) show abduction (opening) and adduction (closing) of the vocal folds. Muscular attachments (posterior cricoarytenoid and thyroarytenoid) to cartilages in the larynx (arytenoid, cricoid, and thyroid) are illustrated to the right. The vocal folds are white bands of specialized vibratory tissue that are necessary for voice production, and the space between the vocal folds is the glottis. b) Laryngeal valving is regulated across the respiratory cycle, with contraction of the posterior cricoarytenoid opening the glottic space during inspiration, and the thyroarytenoid narrowing the glottic space during early expiration (E1). This narrowing increases early-expiratory subglottic pressure and therefore reduces the initial flow of expired gasses. Representative traces of electromyogram (EMG) activity recorded from posterior cricoarytenoid (top) and thyroarytenoid (bottom) muscles during eupnea prior to (blue traces) and 90 minutes after (purple traces) C2 lateral hemisection in cats show a substantial increase in laryngeal inspiratory EMG activity after injury. Traces are waveform averages of the rectified and smoothed (50 ms) EMGs across one minute of stable eupneic activity. c) An unrectified raw trace of the posterior cricoarytenoid EMG shows increased motor drive immediately following C2 lateral hemisection in spontaneously breathing cats (i.v. pentobarbital anesthesia), that was sustained >90 minutes after injury. d) The plotted mean integrated EMG amplitudes illustrate a significant increase from control (pre-injury) in the periods immediately after the injury [*t*(3) = - 5.1, *p* = 0.02] and 90 minutes post-injury [*t*(3) = -6.3, *p* = 0.008; paired t-tests]. To limit the potential confound of cycle-by-cycle variability, the percent change in amplitude was compared to the median peak amplitude during the control period.

Surgical placement of EMGs proceeded as follows: the digastric muscles were blunt dissected away from the surface of the mylohyoid and electrodes were placed medially in the left mylohyoid. A small horizontal incision was made at the rostral end of the right mylohyoid and an elevation of the caudal edge of the incision revealed the geniohyoid muscle. Electrodes were placed 1 cm from the caudal insertion of the geniohyoid muscle. The thyroarytenoid muscle electrodes were inserted through the cricothyroid window into the anterior portion of the vocal folds, which were visually inspected for placement accuracy post-mortem. Minor rotation of the larynx and pharynx counterclockwise revealed the superior laryngeal nerve, which facilitated placement of the thyropharyngeus muscle electrodes. The thyropharyngeus is a fan shaped muscle with the smallest portion attached to the thyroid cartilage; electrodes were placed in the ventral, caudal portion of the muscle overlaying thyroid cartilage within 5 mm of the rostral insertion of the muscle. To place electrodes within the cricopharyngeus muscle, the larynx and pharynx were rotated counterclockwise to reveal the posterior aspect of the larynx. The edge of the cricoid cartilage was located by palpation and electrodes placed in the cricopharyngeus muscle just cranial to the edge. Thyrohyoid muscle electrodes were inserted approximately 5 mm rostral to the attachment to the thyroid cartilage. The esophagus was blunt dissected from the trachea on the contralateral side, and by elevating the trachea, the dorsal side of the larynx was visualized and the posterior cricoarytenoid EMG placed.

A bilateral auricular block with 1% lidocaine was then performed before placing the animal in the stereotaxic frame. Bilateral crural diaphragm EMGs were placed according to the method detailed by Trelease, Sieck and Harper (1982): the 13^th^ rib was manually palpated to locate its junction with the vertebral body, then wire was passed (in a 1.5 in, 22 gauge needle) caudal and lateral to the head of the 13^th^ rib along the sagittal plane (see Figure 2). Post-mortem assessment confirmed placement. The positions of all electrodes were confirmed by visual inspection (both following electrode placement and post-mortem) and by EMG activity patterns during breathing and swallow, as we have previously published (Pitts, Rose et al. 2013, Spearman, Poliacek et al. 2014, Pitts, Rose et al. 2015, Pitts, Poliacek et al. 2018).

**Figure 2.**
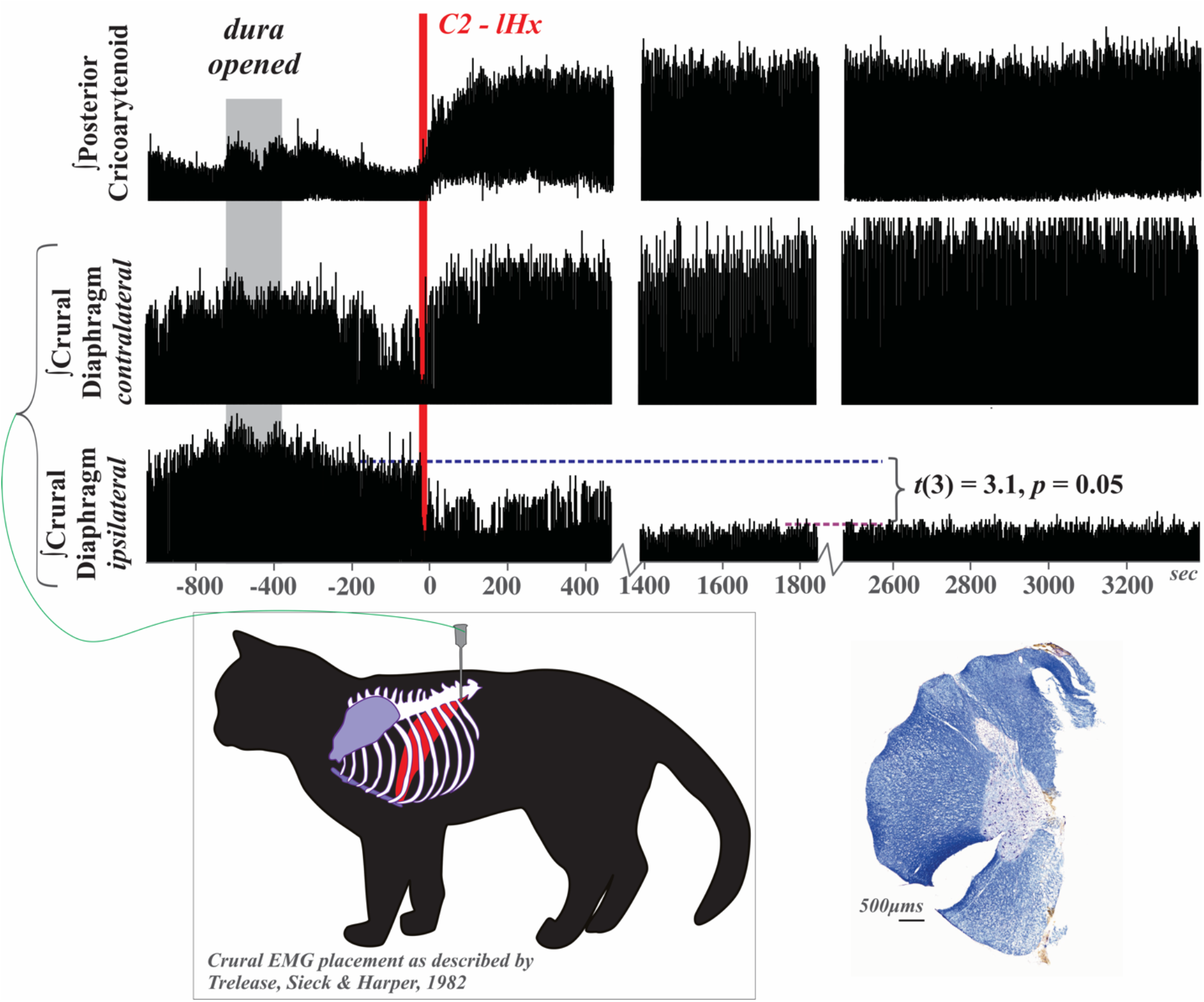
Respiratory drive is altered following cervical spinal cord injury. Electromyogram (EMG) activity was recorded from posterior cricoarytenoid and bilateral crural diaphragm muscles prior to, during, and after C2 lateral hemisection (C2 – *l*Hx) in freely breathing (pentobarbital anesthetized) cats with intact vagi. There was reduction in amplitude, but no evidence of paralysis of the ipsilateral crural diaphragm after lateral hemisection. In 3 of the 4 animals, drive of the contralateral diaphragm increased qualitatively, but the effect was not significant as a group *t*(3) = -1.05, *p* = 0.3. EMG was rectified and smoothed at 50 ms. One minute of stable eupneic breathing before and at 1 hour post-injury was analyzed, and to limit the potential confound of cycle-by-cycle variability, the percent change in amplitude was compared to the median peak amplitude during the control period. Crural diaphragm EMG placement is illustrated (post mortem confirmation example in Figure 3c). A representative example of the C2 lateral hemisection counterstained for Nissl and myelin shows an accurate and complete right side hemisection (left side slit to maintain orientation throughout lesion block during histological processing).

**Figure 3.**
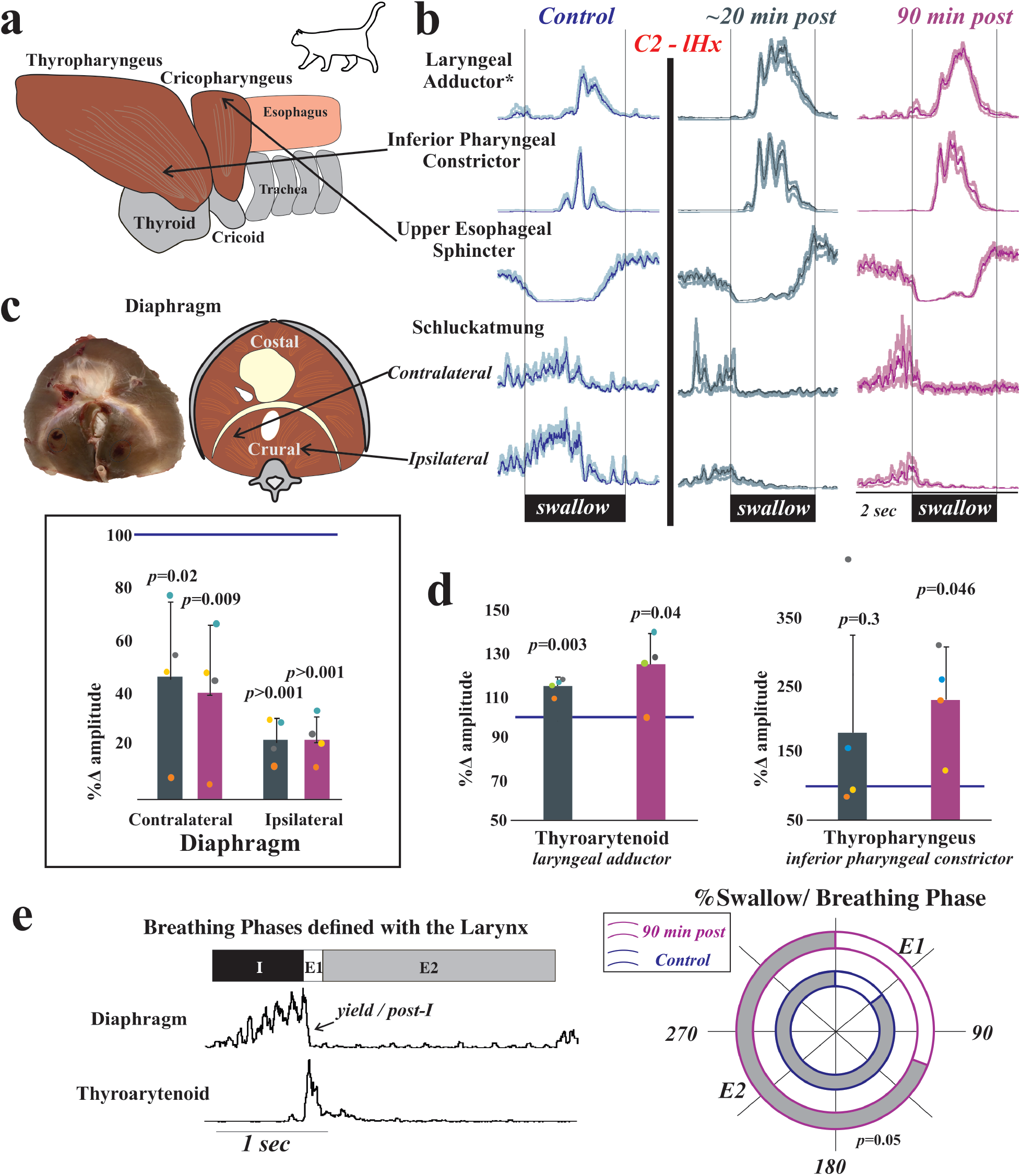
Laryngeal and pharyngeal drive during swallow increases after cervical spinal cord injury. a) While both of the lower pharyngeal muscles (thyropharyngeus and cricopharyngeus) are involved in pharyngeal constriction and maintaining tone in the upper esophageal sphincter, we record from the thyropharyngeus to assess inferior pharyngeal constriction and from the cricopharyngeus to assess upper esophageal sphincter activity. b) Representative traces of electromyogram (EMG) activity recorded from muscles during swallow prior to (blue) and ∼20 minutes (gray) and 90 minutes (purple) after C2 lateral hemisection (C2 – *l*Hx) in cats show changes after injury. The laryngeal adductor is the thyroarytenoid muscle, and traces are waveform averages of the rectified and smoothed (50 ms) EMGs. Swallows were induced with infusion of 3cc of water into the oropharynx. The control waveform (blue) demonstrates the actuation of the diaphragm during swallow (termed “schluckatmung”) during expiration. Oral infusion of water reliably elicits swallow, and EMG amplitudes for laryngeal closure and pharyngeal constriction significantly increased after injury; swallow-related activity of the crural diaphragm significantly decreased on both sides. c) Post-mortem and illustrated images of the diaphragm show locations of the EMG placements. The plot shows decreases in waveform average amplitudes for swallow-related diaphragm activity at ∼20 (gray) and 90 (purple) minutes post-injury as a percent of the control amplitude. Contralateral crural diaphragm was decreased compared to control at ∼20 minutes [*t*(3) = 3.082, *p* = 0.03] and 90 minutes post-injury [*t*(3) = 4.7, *p* = 0.02]; ipsilateral crural diaphragm was also decreased at ∼20 minutes [*t*(3) = 18.5, *p* < 0.001] and 90 minutes post-injury [*t*(3) = 17.6, *p* < 0.001] (paired t-tests). d) Laryngeal adductor (thyroarytenoid) EMG activity during swallow was significantly increased at ∼20 minutes [*t*(3) = -6.9, *p* = 0.003] and 90 minutes post-injury [*t*(3) = -2.1, *p* = 0.04] compared to control. Inferior pharyngeal constrictor (thyropharyngeus) activity during swallow was significantly increased at 90 minutes post-injury [*t*(3) = -3.3, *p* = 0.045]. e) Representative EMG example of breathing are shown with breathing phases defined using laryngeal drive. The percentages of swallow in each phase were plotted across a 180° circle plot (white indicates swallows in E1; grey indicates swallows in E2; inner circle with blue outline indicates early time point; outer circle with pink outline indicates 90 min time point), and a Wilcoxon signed-rank test detected a significant change in swallow breathing coordination, with significantly greater number of swallows occurring during E1 after injury (*Z* = -1.9, *p* = 0.05).

### Rat Model

The animals were initially anesthetized with gaseous isoflurane (1.5-2% with 100% O_2_) while a femoral intravenous (i.v.) cannula was placed for administration of sodium pentobarbital (25 mg/kg, i.v.). Isoflurane was discontinued and supplementary doses of sodium pentobarbital were administered as needed throughout the experiment. A dose of atropine sulfate (0.01mg/kg, i.v.) was given at the beginning of the experiment to reduce secretions from repeated tracheal stimulation. Following administration of atropine sulfate, a tracheostomy was performed. Body temperature was monitored and maintained at 36.5 ± 0.5 °C (Homeothermic Monitor, Harvard Apparatus). Anesthetic level was evaluated by withdrawal reflex of the forelimb and hindlimb and licking in response to oral water administration.

Using the same techniques described above, EMGs of the costal diaphragm and posterior cricoarytenoid were used to evaluate breathing. Additionally, electrodes were placed bilaterally into the pectoralis muscle to record electrocardiogram (ECG) activity, which was used to remove heart artifact from EMG traces.

### Spinal cord injury

Muscle was separated from the dorsal process and laminae to visualize the entire dorsal aspect of the vertebra overlying the intended spinal lesion level along with the rostral most aspect of the adjacent, caudal vertebra. Bilateral laminectomies were performed at either C2 (cat) or T1 (rat) to expose the spinal cord and the dura mater slit longitudinally along the midline of the entire segment. Following visualization of the dorsal columns and dorsal root entry zones, midline was identified and iridectomy scissors used to cut the left half of the spinal cord (cat) or make a complete spinal transection (rat). Lesion completeness was assessed by visualizing lateral and ventral dural surfaces with magnifiers. Further, with complete transection the cut ends pull away from each other leaving a couple mm gap. Gelfoam, saline and bovine thrombin (91-010; BioPharm Laboratories, LLC) were used to control local bleeding. Once hemostasis was achieved, gelfoam was left in the lesion site and it was covered with saline soaked gelfoam and gauze for the remainder of the experiment.

### Histological Processing for the C2 lateral hemisection

Following the collection of terminal electrophysiology data, animals were perfused transcardially with saline (0.9%) followed by 4% buffered paraformaldehyde (pH 7.4). Spinal cords were removed, lesion level verified, and tissue blocked. Tissue was post-fixed for 24-48 hours, then cryoprotected in 30% buffered sucrose (pH 7.4) and sectioned on a cryostat (25 μm) at -20-22°C. Every 40th section (every 1,000 μm) of the lesion block was processed with Nissl (Cresyl Violet Acetate, Sigma) and myelin (Eriochrome Cyanine R, Merck) stains to allow identification of the injury epicenter and evaluation of the extent of the lesion (Mondello, Jefferson et al. 2015). Sections were dehydrated through serial ethanols (70 to 100%) and coverslipped out of xylenes (Fisher) using DPX mounting media (Electron Microscopy Sciences), and photographed using a Zeiss Imager. Z2 and Stereo Investigator (MicroBrighfield).

## Results

First, we assessed the effect of C2 lateral hemisection in cats on laryngeal drive during breathing. Figure 1 demonstrates an increase in posterior cricoarytenoid EMG activity during inspiration (percent change ∼20 minutes post-injury 189 ± 35, 90 minutes post-injury 173 ± 23). The posterior cricoarytenoid attaches from the cricoid to the arytenoid cartilages, and its contraction rotates the arytenoid cartilage to move the vocal folds into an abducted (open) position. Increased amplitude represents an increased glottic space and decreased laryngeal resistance during inspiration.

Second, we assessed the effect of C2 lateral hemisection on bilateral crural diaphragm activity during breathing. Ipsilateral to the injury, the crural diaphragm EMG was significantly reduced (Figure 2), but there was not a significant increase in contralateral recruitment. Obtaining reliable bilateral diaphragm recordings was essential to the assessment, but opening the abdominal cavity affects chest wall activity (Beecher 1933, Beecher 1933, Farkas and De Troyer 1989, Mondal, Abu-Hasan et al. 2016) and alters swallow breathing coordination (Pitts, Rose et al. 2015). Therefore, we instead used the method described by Trelease, et al. (1982), (Figure 2) which produced stable bilateral diaphragm recordings while maintaining normal respiratory mechanics. There was a significant depression of ipsilateral crural diaphragm EMG activity after C2 lateral hemisection (62 ± 25 percent change post-injury), but no animal had a complete termination of all activity. Additionally, 75% of animals had an increase in contralateral diaphragm recruitment, but this did not reach significance. The smallest contralateral effect of injury was at 90% of control, and the largest effect was at 174% of control, which is displayed in Figure 2. Post-mortem tissue examination confirmed that all animals had complete lateral hemisections at C2 and confirmed correct electrode placement in the crural diaphragm.

Third, we determined the effect of C2 lateral hemisection on regulation of the upper airway during swallow. Measurements from the full complement of muscles recorded are reported in supplemental Figure S1. These muscles were chosen because they are vital to the operation of the aerodigestive dual valve mechanism (Pitts, Rose et al. 2013). In this scenario, the upper esophageal sphincter and larynx must coordinate their movements to control the flow of air/liquid from the upper airway into either the trachea or esophagus in the thoracic cavity. The primary outcome of the injury was that the thyroarytenoid’s EMG activity increased acutely (percent change 113 ± 6, ∼20 minutes post-injury,) and this increase was maintained (90 minutes post-injury, 118 ± 20). The thyroarytenoid spans the larynx and constitutes the bulk of the vocal folds. Increased thyroarytenoid contraction ensures that the glottic space is closed during swallow, reducing aspiration risk. However, thyropharyngeus activity (operating as the inferior pharyngeal constrictor) also increased after injury (percent change 90 minutes post-injury 229 ± 79). The thyropharyngeus is a fan-shaped muscle that attaches to the lateral aspect on the thyroid cartilage and constitutes the bulk of the pharyngeal wall. Increased thyropharyngeus activity during swallow adds to the pressure exerted on the laryngeal orifice, but is also necessary to maintain a clear pharynx for the subsequent breath. There was also no significant change in the number of swallows stimulated per trial [control 3.5 ± 0.6, ∼20 min post-injury 4.7 ± 2.7 (*p =* 0.5), and 90 minutes post-injury 3 ± 0.7 (*p =* 0.8)].

Fourth, we found that diaphragm activity during swallow was bilaterally suppressed after lateral C2 hemisection (percent change 90 minutes post-injury, 22 ± 9 ipsilateral; 40 ± 25 contralateral). This finding was not anticipated, given that the C2 lateral hemisection only suppressed ipsilateral diaphragm breathing activity, with 3 out of 4 animals showing increased contralateral EMG activity during breathing.

Fifth, there was a significant change in swallow-breathing coordination. Using the defined breathing phases described in Figure 2e, each swallow initiation was marked in the corresponding phase of breathing. Following spinal cord injury, there was a significant increase in the percentage of swallows that occurred during E1 (i.e. post-I or yield phase).

Finally, we wanted to determine if our observations in the cat were relevant in another animal model, and to determine if the chest wall is a major source of feedback for laryngeal regulation. Figure 4 illustrates two features of data from rat T1 transections that were similar to the cat data after injury: 1) increased laryngeal inspiratory activity, and 2) increased diaphragm activity during breathing. A necessary component of the rat experiments was reliable recording of the posterior cricoarytenoid muscle, which is small and very close to the pharyngeal wall. We used the same EMG technique as above, but placed one wire in each of the bilateral muscle bellies. As in the cat C2 lateral hemisection, an increase in inspiratory laryngeal activity (170 ± 34 percent change) was the first feature noted after rat T1 complete spinal cord transection, and this effect was stable over the recording period. We recorded costal (midline) diaphragm activity in the rat experimental preparations, rather than crural (bilateral) activity, because sided effects were not necessary to the analysis. In contrast to the cat C2 lateral hemisection, diaphragmatic respiratory drive increased after T1 transection in every rat (215 ± 63 percent change), and this effect was significant (Figure 4).

**Figure 4.**
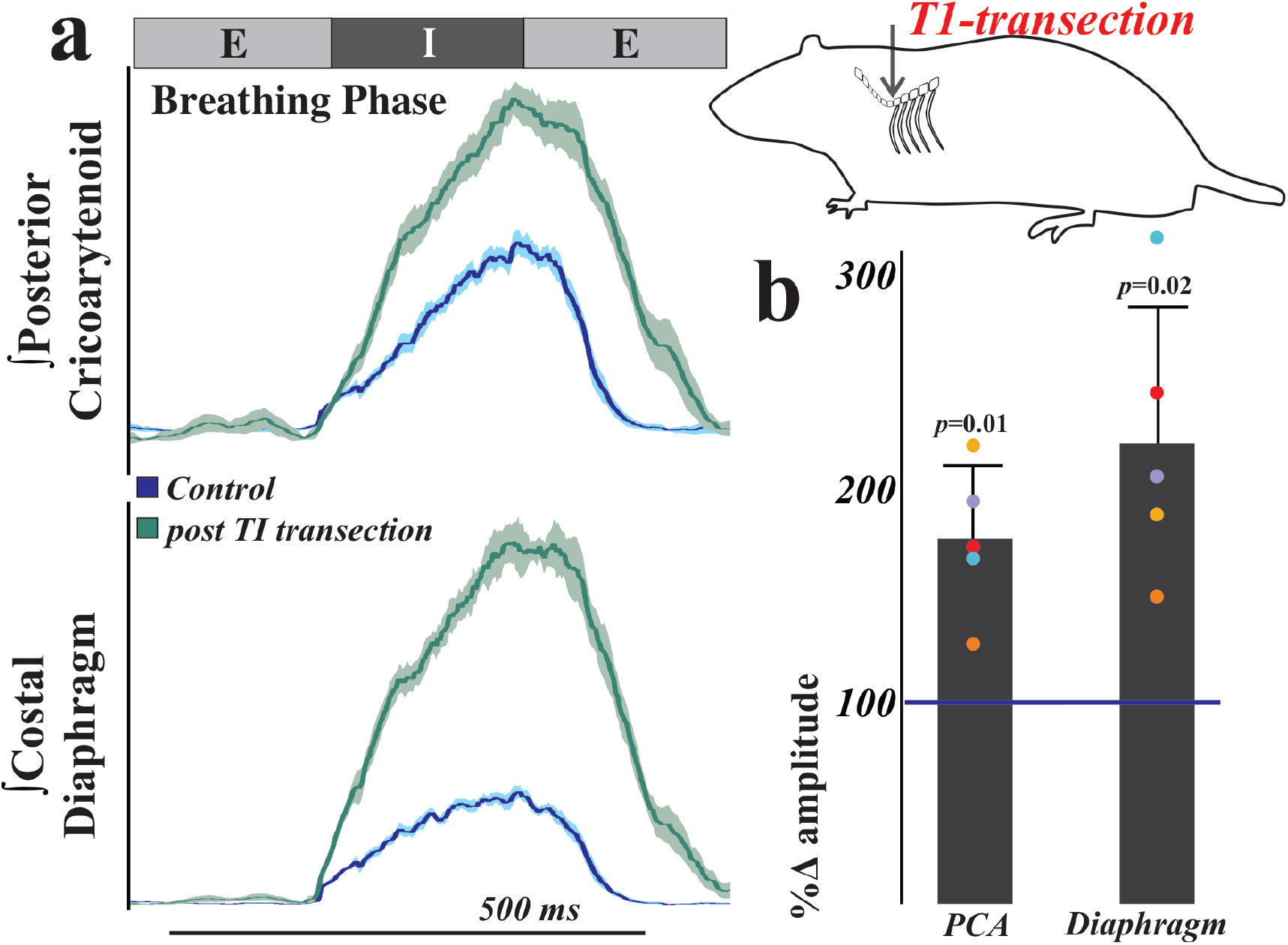
Respiratory drive is altered after T1 spinal cord transection in the rat. To test the hypothesis that thoracic afferent feedback may be a significant contributor to laryngeal regulation, we performed T1 total transections in freely breathing (vagi intact) pentobarbital anesthetized female Sprague Dawley rats. A) Representative traces of electromyogram (EMG) activity recorded from posterior cricoarytenoid (top) and costal diaphragm (bottom) muscles during eupnea prior to (blue traces) and ∼20 minutes after (green traces) total transection at the T1 spinal level in rats show increased activity after injury. Traces are waveform averages of the rectified and smoothed (50 ms) EMGs from 1 minute of stable eupneic activity. B) EMG amplitudes 20 minutes after transection were compared to the median amplitude during the control period to control for cycle-by-cycle variability and plotted as a percent of control. As assessed by paired t-tests, there were significant increases in posterior cricoarytenoid [*t*(4) = -4.6, *p* = 0.01] and costal diaphragm [*t*(4) = -4.04, *p* = 0.02] amplitudes post-transection.

## Discussion

Swallow is a primitive and critical behavior necessary for nutrition and survival (Negus 1942, Doty and Bosma 1956, Dullemeijer 1956, Gupta 1971, Herring and Scapino 1973, Miller 1982). Some aspects of the swallow motor pattern changed as vertebrates evolved complexity, and it is a common point of disability across a range of human neuro-developmental, traumatic, and degenerative diseases. As the larynx is the central point in regulation of the upper airway, it is vital to regulating airway protective risk. This study demonstrated cross-species effects of spinal cord injury on the upper airway that are indicative of increased risk for aspiration. Moreover, there was behavior-specific facilitation of upper airway motoneurons contained only with the brainstem. Additionally, while we predicted that C2 lateral hemisection would decrease ipsilateral crural diaphragm activity during breathing and swallowing, there was a bilateral suppression during swallow, demonstrating that swallow may be more susceptible to disorder following spinal cord injury.

### Intrinsic laryngeal muscles

The posterior cricoarytenoid is the intrinsic laryngeal muscle that regulates abduction of the vocal folds during inspiration; it is innervated by the recurrent laryngeal nerve. In cat, its innervating motoneurons are reported throughout the nucleus ambiguus, with the largest neuron size in the caudal portion (Yoshida, Miyazaki et al. 1982, Pásaro, Lobera et al. 1983). In rat, multipolar motoneurons are described in the compact and loose formations of the nucleus ambiguus, mainly in the middle portion of the column Berkowitz et al (1999), and they have an augmenting discharge pattern during inspiration that is similar to the phrenic nerve output (Berkowitz, Chalmers et al. 1999) (Berkowitz, Chalmers et al. 1999). These motoneurons have extensive excitatory and inhibitory synaptic terminals (Hayakawa, Zheng et al. 1999), and their connections are found throughout the respiratory and swallow regulatory neuraxis (Waldbaum, Hadziefendic et al. 2001). Innervating medullary areas include: several regions of the nucleus tractus solitarius; the midline reticular formation (including the medullary raphe); the pre-Bötzinger complex; and gigantocellular and lateral paragigantocellular reticular nuclei. Innervating pontine areas include: the Kolliker-Fuse nucleus, locus coeruleus, sub-coeruleus, lateral tegmental field, the A5 catecholaminergic cell group, and the medial pontine reticular formation (Waldbaum, Hadziefendic et al. 2001).

In contrast, the thyroarytenoid is a primary laryngeal adductor that constitutes the bulk of the vocal folds; it is also innervated by the recurrent laryngeal nerve. In cat, its innervating motoneurons are found in the caudal two-thirds of the nucleus ambiguus continuing past obex (Portillo and Pásaro 1988), and are dorsal to the location of the posterior cricoarytenoid motoneurons (Yoshida, Miyazaki et al. 1982). They appear as loose group with a consistent relatively large size (Pásaro, Lobera et al. 1983). In the rat, they are found primarily in the medial nucleus ambiguus in its caudal portion (Hernández-Morato, Valderrama-Canales et al. 2013). In a pseudorabies tracing study, Van Daele and Cassell (2009) found thyroarytenoid connections in areas similar areas to those reported for the posterior cricoarytenoid, with some additional areas. These additional connections were labeled in more regions of the nucleus tractus solitarius and reticular formation (including intermediate and parvocellular formations); the periaqueductal gray; along the ventral respiratory group including the Bötzinger Complex; and in the phrenic nucleus.

While many brainstem connections to the motoneurons for these muscles have been described, much less is known about the potential anatomical pathways connecting the larynx and spinal cord. In Remmers’ 1973 papers, stimulation of external intercostal nerve afferents evoked laryngeal reflexes that were eliminated by an acute lateral cut (lateral to the ventral horn) at C3. When the stimulation was applied during inspiration, the posterior cricoarytenoid was inhibited, but when stimulated during expiration, the posterior cricoarytenoid and thyroarytenoid were both excited (Remmers 1973, Remmers and Tsiaras 1973). Our use of the T1 transection in the rat underscores the hypothesis that the increase in the posterior cricoarytenoid after injury may be due to loss of afferent feedback from the chest wall. Information from these afferents travels through several tracts to the brainstem, but the lateral area lesioned by Remmers would have included the spinocerebellar, lateral spinothalamic, and spinoreticular tracts. This is also supported by work from Kirkwood and colleagues showing increased excitation (especially during the E1/post-I phase) of respiratory motoneurons above the lesion in thoracic hemisections which spared the dorsal columns (Ford, Anissimova et al. 2016). Our recent works studying the effects of acute cerebellectomy on breathing, swallow, and cough suggest that the ascending cerebellar pathways are not responsible (Reed, English et al. 2019, Musselwhite, Shen et al. 2021), leaving the lateral spinothalamic and spinoreticular tracts as possible key pathways.

### Dual Valve Hypothesis

A very old hypothesis posits that the diaphragm is active during swallow (Rosenthal 1861, Bidder 1865, Blumberg 1865, Waller and Prevost 1869, Kronecker and Meltzer 1880, Kronecker and Meltzer 1880, Kronecker and Meltzer 1881, Meltzer 1882). The diaphragm is accepted as the major source of negative pressure which draws air into the lungs during breathing. Our data are consistent with the old but recently-neglected hypothesis that the diaphragm acts in a similar aspiration pump role during swallow to draw food/liquid into the esophagus. A reasonable question is, “How can the diaphragm be active during swallow without causing aspiration of material into the lungs?” As published in Pitts, et al (2013), the larynx has classically been depicted as a valve. We extended this view to a dual valve system to incorporate the fact that the upper esophageal sphincter works cooperatively with the laryngeal muscles that control the opening to the trachea. In this way, only one “valve” is open at any one time, which ensures effective mechanical movement of air/bolus into the thoracic cavity without a “leak” from the cooperating valve. Common to all behaviors, the diaphragm would function by using negative pressure to draw into the thoracic cavity, including breathing, cough, sneeze, and swallow. Unlike breathing, however, swallow is aided by cooperation from other structures after the bolus passes. This includes closure of the oral cavity and contraction of the tongue and upper pharyngeal muscles, which produces a positive pressure above the food/liquid bolus (McConnel, Cerenko et al. 1988, McConnel, Cerenko et al. 1988, McConnel, Hester et al. 1988).

While the upper airway is able to exert powerful force on the bolus, dysphagia is common among patients with disordered negative intra-thoracic pressure during swallow (McConnel, Mendelsohn et al. 1986, McConnel, Mendelsohn et al. 1987, Mendelsohn and McConnel 1987, McConnel 1988, McConnel, Cerenko et al. 1988, McConnel, Cerenko et al. 1988, McConnel, Cerenko et al. 1988, McConnel, Hester et al. 1988, Cerenko, McConnel et al. 1989, Ku, Ma et al. 1990, McConnel, Guffin et al. 1991, McConnel, Guffin et al. 1992). There is extensive literature on how cSCI affects breathing-related phrenic nerve and diaphragm recruitment from the groups of Goshgarian (Hadley, Walker et al. 1999, Goshgarian 2009), Sieck and Mantilla (Mantilla and Sieck 2003, Mantilla and Sieck 2011, Mantilla, Bailey et al. 2012),, Mitchell and Fuller (Fuller, Johnson et al. 2002, Fuller, Johnson et al. 2003, Mitchell and Johnson 2003), Reier and Lane (Lane, Fuller et al. 2008, Lane, Lee et al. 2009), Alilain and Silver (Alilain, Li et al. 2008, Alilain and Silver 2009, Alilain, Horn et al. 2011, Sharma, Alilain et al. 2012, Awad, Warren et al. 2013), and others (Ginsborg and Hirst 1972, Golder, Reier et al. 2001, Golder, Fuller et al. 2003, Polentes, Stamegna et al. 2004, Baussart, Stamegna et al. 2006, DiMarco 2009); however, we have little knowledge about the effects of cSCI on swallow. The present data demonstrate a bilateral reduction of crural diaphragm activity during swallow, with only an ipsilateral reduction during breathing following C2 lateral hemisection. This provides further evidence of behavior-specific effects, and implies that swallow may be more affected by cervical spinal cord injury than breathing.

We also found that spinal cord injury produced a significant change in swallow breathing coordination, with more post-injury swallows occurring during early expiration (in the E1 phase; also termed post-I or yield). The Kolliker-Fuse nucleus in the pons has been proposed to exert firm control over swallow-breathing coordination, as well as post-I laryngeal adduction (Dutschmann and Herbert 2006, Bautista and Dutschmann 2014). However, we observed a permissive effect of swallows occurring during E1, with a behavior-specific change in laryngeal adductor activity after a lesion to the spinal cord rather than the pons. There is a difference in the methodological procedures, as we present EMG activity recorded selectively from the thyroarytenoid muscle, whereas the Dutschmann studies recorded a collection of action potentials from the cervical vagus nerve. However, human studies strongly support pontine modulation of swallow motor activity based on the clinical utility of activating trigeminal nerve c-fibers before/during eating in patients with dysphagia (Nakato, Manabe et al. 2017, Wang, Wu et al. 2019, Cui, Yin et al. 2020). Collectively, these findings along with ours strongly suggest that both the spinal cord and pons contribute to swallow breathing coordination, and their precise roles are yet to be determined. Based upon our identification of spinal contributions, swallow likely modulates breathing in a complex rather than straightforward manner, involving multiple central regions.

Dysphagia, especially aspiration of oral secretions, is a significant risk factor for community acquired pneumonia (Almirall, Rofes et al. 2013) and hospitalization (Komiya, Ishii et al. 2015). The 2016 annual statistical report from the Spinal Cord Injury Model Systems project says that cervical SCI accounts for 54.2% of the total population, with high cervical (C1-C4) injury representing 21%. In patients for whom cause of death was known, 65.3% of deaths were due to pneumonia, thus it is the leading known cause. Due to the wide array of functions that involve laryngeal regulation, its dysfunction is of particular concern. The large and unexpected upper airway response to cervical and thoracic lesions demonstrates a need for future key studies investigating the connections between the spinal cord and the larynx. Additionally, the behavior-specific effects of the lesions necessitate comprehensive studies to predict the clinical effects of spinal cord injury in humans. There is a strong need for novel therapies or pharmaceutical developments to help these patients at risk for upper airway dysfunction.

