## Supplemental Data for "Laryngeal and swallow dysregulation following acute cervical spinal cord injury"

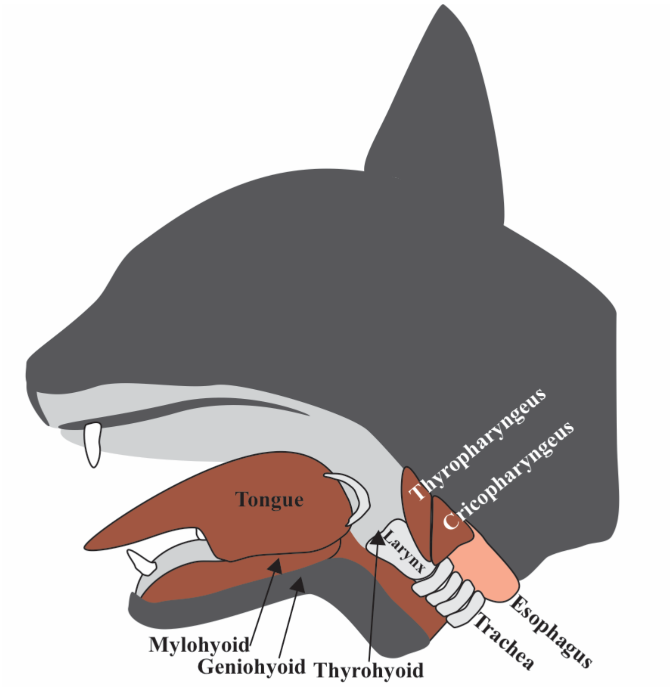


| | ~20 min post | | | | 90 min post | | | |
|---|---|---|---|---|---|---|---|---|
| | **Mean** | **±** | **SD** | ***p-value*** | **Mean** | **±** | **SD** | ***p-value*** |
| **Mylohyoid** | 157 | ± | 80 | 0.12 | 161 | ± | 44 | *0.03* |
| **Geniohyoid** | 357 | ± | 449 | 0.21 | 204 | ± | 121 | 0.14 |
| **Thyrohyoid** | 132 | ± | 14 | *0.03* | 203 | ± | 44 | *0.03* |
| **Cricopharyngeus** | 133 | ± | 52 | 0.19 | 143 | ± | 8 | *0.01* |

Figure S1. Illustration showing supplemental muscles for swallow EMG placement. There was a significant increase in EMG amplitude at ~20 minutes post-injury for mylohyoid, thyrohyoid and the cricopharyngeus (acting as the upper esophageal sphincter). The EMG for the geniohyoid was placed on the side contralateral to the spinal cord injury, while the EMGs for the other muscles were ipsilateral to the injury. This was standard practice from our previous experiments because of anatomical constraints, but is a limitation to the current design.

Table S1. Cardiorespiratory measures before and after cSCI. No animals were placed on a ventilator or given supplemental oxygen.

| | Pre-injury | | | Post-injury | | | F-statistic | *p*-value |
|---|---|---|---|---|---|---|---|---|
| | **Mean** | **±** | **SD** | **Mean** | **±** | **SD** | | |
| **Heart Rate** | 195 | ± | 10 | 163 | ± | 5 | 21.2 | *0.004* |
| **BP- Systolic** | 145 | ± | 26 | 133 | ± | 17 | 0.4 | 0.2 |
| **BP- Diastolic** | 122 | ± | 15 | 108 | ± | 16 | 1.2 | 0.2 |
| **end-tidal $CO_2$** | 31 | ± | 7 | 32 | ± | 19 | 0.1 | 0.6 |
| **Temperature** | 35 | ± | 1 | 35 | ± | 11 | 0.02 | 0.5 |
| **Respiratory Rate** | 33 | ± | 11 | 28 | ± | 10 | 0.5 | 0.3 |
| **Fi-$O_2$** | 21 | ± | 0 | 21 | ± | 0 | - | - |
| **p$CO_2$** | 26 | ± | 7 | 30 | ± | 123 | 0.5 | 0.1 |
| **p$O_2$** | 106 | ± | 10 | 110 | ± | 139 | 0.5 | 0.3 |
| **pH** | 7.4 | ± | 0.1 | 7.4 | ± | 0.6 | 0.5 | 0.7 |
